# Molecular mapping of *qBK1*^*Z*^, a major QTL for bakanae disease resistance in rice

**DOI:** 10.1101/2020.05.18.101725

**Authors:** Sais-Beul Lee, Namgyu Kim, Sumin Jo, Yeon-Jae Hur, Ji-youn Lee, Jun-Hyeon Cho, Jong-Hee Lee, Ju-Won Kang, You-Chun Song, Maurene Bombay, Sung-Ryul Kim, Jungkwan Lee, Young-Su Seo, Jong-Min Ko, Dong-Soo Park

## Abstract

Bakanae disease is a fungal disease of rice (*Oryza sativa* L.) caused by the pathogen *Gibberella fujikuroi* (also known as *Fusarium fujikuroi*). Recently the disease incidence has increased in several Asian countries and continues to spread throughout the world. No rice varieties have been developed yet to be completely resistant to this disease. With increasing need to identify various genetic resources to impart resistance to local elite varieties, this study was carried out to identify novel quantitative trait loci (QTLs) from an *indica* variety Zenith. We performed a QTL mapping using 180 F_2:9_ recombinant inbred lines (RILs) derived from a cross between the resistant variety, Zenith, and the susceptible variety, Ilpum. A primary QTL study using the genotypes and phenotypes of the RILs indicated that the locus *qBK1*^*z*^ conferring bakanae disease resistance from the Zenith was located in a 2.8 Mb region bordered by the two SSR markers, RM1331 and RM3530 on chromosome 1.

The log of odds (LOD) score of *qBK1*^*z*^ was 13.43, accounting for 30.9% of the total phenotypic variation. A finer localization of *qBK1*^*z*^ was delimited at an approximate 730 kb interval in the physical map between Chr01_1435908 (1.43 Mbp) and RM10116 (2.16 Mbp). The development of a rice variety with a higher level of resistance against bakanae disease is a major challenge in many rice growing countries. Introducing *qBK1*^*z*^ or pyramiding with other previously identified QTLs could provide effective genetic control of bakanae disease in rice.

## Introduction

Bakanae disease, which means foolish seedling in Japanese, was firstly identified in 1828 in Japan [1], and has widely distributed in temperate zone as well as tropical environment and occurring throughout rice growing regions of the world [2].

Four *Fusarium* species including *F. andiyazi, F. fujikuroi, F. proliferatum* and *F. verticillioides* in the *G. fujikuroi* species complex have been associated with Bakanae disease in rice [3]. This disease typically seed-borne fungus, but may occur when the pathogen is present in plant material or soil. Infected seeds/plants result in secondary infections [4], which spreads through wind or water that carries the fungal spores from one plant to another. Bakanae disease has different symptoms starting from pre-emergence seedling death to grain infection at maturity such as tall, lanky tillers with pale green flag leaves. Infected plants have fewer tillers, and plants surviving till maturity bear only empty panicles [5], resulting in yield loss [6, 7]. Low plant survival and high spikelet sterility [5] may account for yield losses up to 50% in Japan [6, 7], 3.0-95% in India [2, 8, 9], 3.7-14.7% in Thailand, 5-23% in Spain, 40% in Nepal [10], 6.7-58.0% in Pakistan [11], 75% in Iran [12], and to 28.8% in Korea [13]. Hot water immersion and fungicide treatment are most common ways for seed disinfection [2, 14, 15]. However, both the hot water treatment and application of fungicide are insufficient to control bakanae disease. Thermal effect does not reach the pericarp of the severely infected rice seeds. The application of fungicides is not functioning well for destroying the spores of this fungal pathogen, and some pathogen showed resistance to the fungicides [13, 16-18]. Therefore, the genetic improvement of rice using the QTLs/genes providing the bakane disease resistance would be a more effective way to control bakanae disease.

Several QTLs associated with bakanae disease resistance have been identified and those can be used for marker-assisted selection in rice breeding as well as for understanding the mechanisms of resistance. Yang et al. [19] identified two QTLs located on chromosome 1 and chromosome 10 by *in vitro* evaluation of the Chunjiang 06/TN1 doubled haploid population. Hur et al. [20] identified a major QTL, *qBK1*, on chromosome 1 from 168 BC_6_F_4_ near isogenic lines (NILS) generated by crossing the resistant *indica* variety Shingwang with susceptible *japonica* variety Ilpum. Lee et al. [21] delimited the location of *qBK1* to 35 kb interval between two InDel markers, InDel 18 (23.637 Mbp) and InDel 19-14 (23.672 Mbp). Fiyaz et al. [9] identified three QTLs (*qBK1*.*1, qBK1*.*2*, and *qBK1*.*3*) on chromosome 1 and one QTL (*qBK3*.*1*) on chromosome 3 from the highly resistant variety Pusa 1342. Physical location of *qBK1*.*1* is close to *qBK1* [21]. Ji et al. [22] mapped a major QTL (*qFfR1*) in 22.56–24.10 Mbp region on chromosome 1 using 180 F_2:3_ lines derived from a cross between a resistant Korean *japonica* variety Nampyeong and a susceptible Korean *japonica* line, DongjinAD. Volante et al. [23] identified *qBK1_628091* (0.6 Mbp to 1.0 Mbp on chromosome 1) and *qBK4_31750955* (31.1 Mbp to 31.7 Mbp on chromosome 4) by GWAS (Genome Wide Association Study) approach using 138 *japonica* rice germplasm. Kang et al [24] discovered the QTL *qFfR9* at 30.1 centimorgan (cM) on chromosome 9 from a *japonica* variety Samgwang. Lee et al. [15] found the QTL *qBK1*^*WD*^ located between markers chr01_13542347 (13.54 Mb) and chr01_15132528 (15.13 Mb) from the *japonica* variety Wonseadaesoo. They also found that resistance of gene pyramided lines harboring two QTLs, *qBK1*^*WD*^ and *qBK1* was significantly higher than those with only *qBK1*^*WD*^ or *qBK1*. This finding relates to the problem of varieties with single resistance gene losing its effect against new populations of fungal isolates [25]. Hence, identifying new resistance genes from diverse sources is important for rice breeding programs to acquire durable resistance against bakanae disease by enhancing the resistance level and/or help to overcome the breakdown of resistance genes.

In this study, we aimed to provide a new genetic source, *qBK1*^*z*^ with detailed gene locus information for developing resistant rice lines which contains single or multiple major QTLs to enhance bakanae disease resistance.

## Materials and Methods

### Plant materials

Zenith, a medium grain type *indica* variety of USA released in 1936, was identified as resistant to bakanae disease in a preliminary screening of rice germplasm (data not shown) using the large-scale screening method developed by Kim et al. [26]. We generated 180 F_2:9_ RILs from a cross between susceptible variety, Ilpum, and resistant variety, Zenith for QTL analysis. The population was developed in the experimental fields of the National Institute of Crop Science of the Rural Developmental Administration in Miryang, Korea.

### Evaluation of bakanae resistance

The inoculation and evaluation of bakanae disease were conducted using a method that described by Lee et al. [21]. The isolate CF283 of *F. fujikuroi* (imperfect stage of *G. fujikuroi*) were obtained from National Academy Agricultural Science in Korea. Isolate was inoculated in potato dextrose broth (PDB) and cultured at 26 °C under continuous light for one week. The *F. fujikuroi* culture was washed by centrifugation with distilled water. Regarding the standardization of the density of the homogenized inoculum, the fungal spore concentration was adjusted to 1 × 10^6^ spores/mL with a hemocytometer. Forty seeds per each line were placed in a tissue-embedding cassette (M512, Simport, Beloeil, QC, Canada). Before inoculation, the seeds in the in the tissue-embedding cassette were surface sterilized with hot water (57 °C) for 13 min, then allowed to drain and cool. Subsequently, the seeds were soaked in the spore suspensions (1 × 10^6^ spores/mL) for 3 days for inoculation with gentle shaking four times a day for equilibration. After inoculation, thirty seeds per line were sown in commercial seedling tray, and seedlings were grown in a greenhouse (28 ± 3 °C day/23 ± 3 °C night, 12 h light). Bakanae disease symptom on each line was evaluated by calculating the proportion of healthy plants at 1 month after sowing. The healthy and non-healthy plants are classified as described by Kim et al. [26]. The plants exhibit elongation with thin and yellowish-green, elongation then stunted growth, stunted growth, dead seedling were classified as non-healthy plants. The plants showing same phenotype with untreated plants, slight elongation then normal growth without thin and yellowish-green was regarded as healthy plants.

### Localization of *F. fujikuroi* in Zenith and Ilpum plants

Shoot base of 10 day-old seedlings derived from Zenith and Ilpum seeds inoculated by GFP-tagged *F. fujikuroi* isolate CF283 [15] was observed under confocal laser-scanning microscopy (LSM-800, Zeiss, Germany) at GFP channel, and images were obtained using Zeiss LSM Image Browser. All experiments were conducted twice with at least three replicates.

### DNA extraction and polymerase chain reaction

Genomic DNA from young leaf tissue was prepared according to the CTAB method [27] with minor modifications. Polymerase chain reaction (PCR) was performed in 25-µL reaction mixture containing 25 ng template DNA, 10 pmol of each primer, 10× e-Taq reaction buffer, 25 mM MgCl2, 10 mM dNTP mix and 0.02 U of SolGent™ e-Taq DNA polymerase (SolGent, Daejeon, South Korea). The reaction conditions were set as follows: initial denaturation at 94°C for 2 min; 35 cycles of denaturation at 94°C for 20 s, annealing at 57°C for 40 s and extension at 72°C for 40 s; and a final extension at 72°C for 7 min. The amplification products were electrophoresed on a 3% (w/v) agarose gel and visualized by ethidium bromide staining.

### QTL analysis of the F_2:9_ population and development of InDel markers for fine mapping

Polymorphic SSR markers (n = 164) that were evenly distributed on rice chromosomes were selected from the Gramene database (http://www.gramene.org). These markers were used to construct a linkage map and for QTL analysis of the F_2:9_ populations. The linkage map was constructed using Mapmaker/Exp v.3.0, and the genetic distance was obtained using the Kosambi map function [28]. Putative QTLs were detected using the composite interval mapping (CIM) function in WinQTLcart v.2.5 (WinQTL cartographer software [29]), with the threshold being set based on the results of 1000 permutation tests. A logarithm of the odds (LOD) ratio threshold of 3.0 was used to confirm the significance of a putative QTL. InDel markers were developed based on the fragment size differences in the sequence (in the range of 20 bp) between *japonica* (Gramene database http://www.gramene-.org) and *indica* (BGI-RIS; http://rice.genomics.org.cn) in the target region on chromosome 1. The primers were designed using Primer3 software (http://web.-bioneer.co.kr/cgi-bin/primer/primer3.cgi).

### Statistical analysis

Statistical differences between means were analyzed using Duncan’s multiple range test after one-way analysis of variance (ANOVA). The level of significance was designated as P < 0.05 and was determined using the SAS Enterprise Guide 4.3 program (SAS Institute Inc., Cary, NC, USA).

## Results

### Bakanae disease bioassay in parents and F_2:9_ RILs

The proportion of healthy Zenith (resistant) and Ilpum (susceptible) plants was investigated after inoculation of the virulent *F. fujikuroi* isolate CF283 [26]. Most of Zenith plants did not exhibit a thin and yellowish-green phenotype, which is a typical symptom of bakanae disease as unlike Ilpum (Fig. 1A). The proportion of healthy plants was significantly different between Ilpum (14.3%) and Zenith (63.2%) (Fig. 1B).

**Fig 1.**
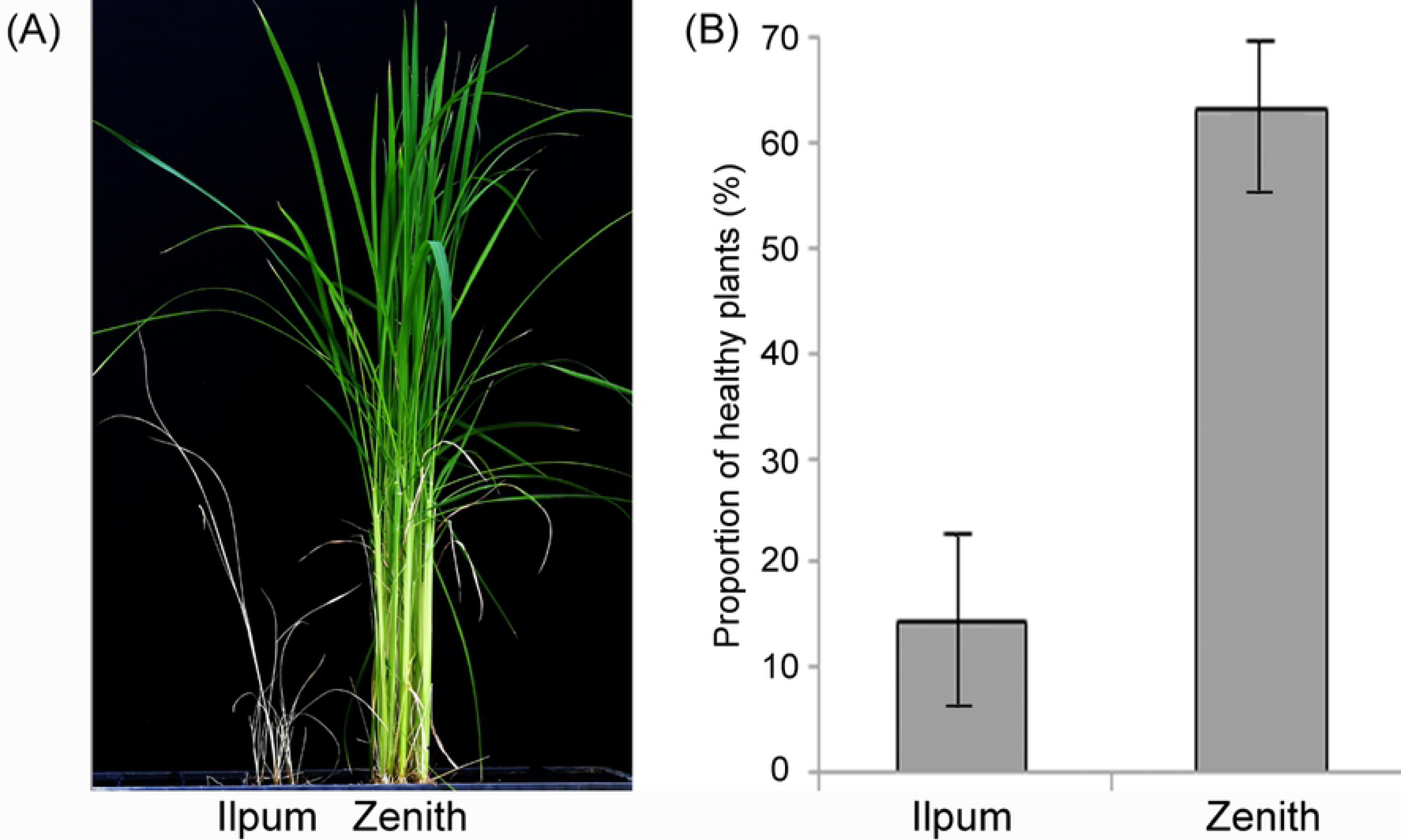
Bioassay of parents (Ilpum and Zenith) against *F. fujikuroi* isolate CF283. (A) Plant phenotype from the pathogen-inoculated seeds of Ilupm and Zenith. (B) Portion of healthy plants from the pathogen-treated seeds.

Zenith and Ilpum were inoculated with green fluorescent protein (GFP)-tagged *F. fujikuroi* isolate CF283. Ten days after inoculation, plants with typical disease symptom of each variety were subjected to a confocal microscopy analysis. Confocal imaging of radial and longitudinal sections of the basal stem showed that the fungus penetrated and was localized easily and abundantly at vascular bundle, mesophyll tissue and hypodermis in the susceptible Ilpum variety while only background level of GPF signal was detected in the resistant Zenith (Fig. 2).

**Fig 2.**
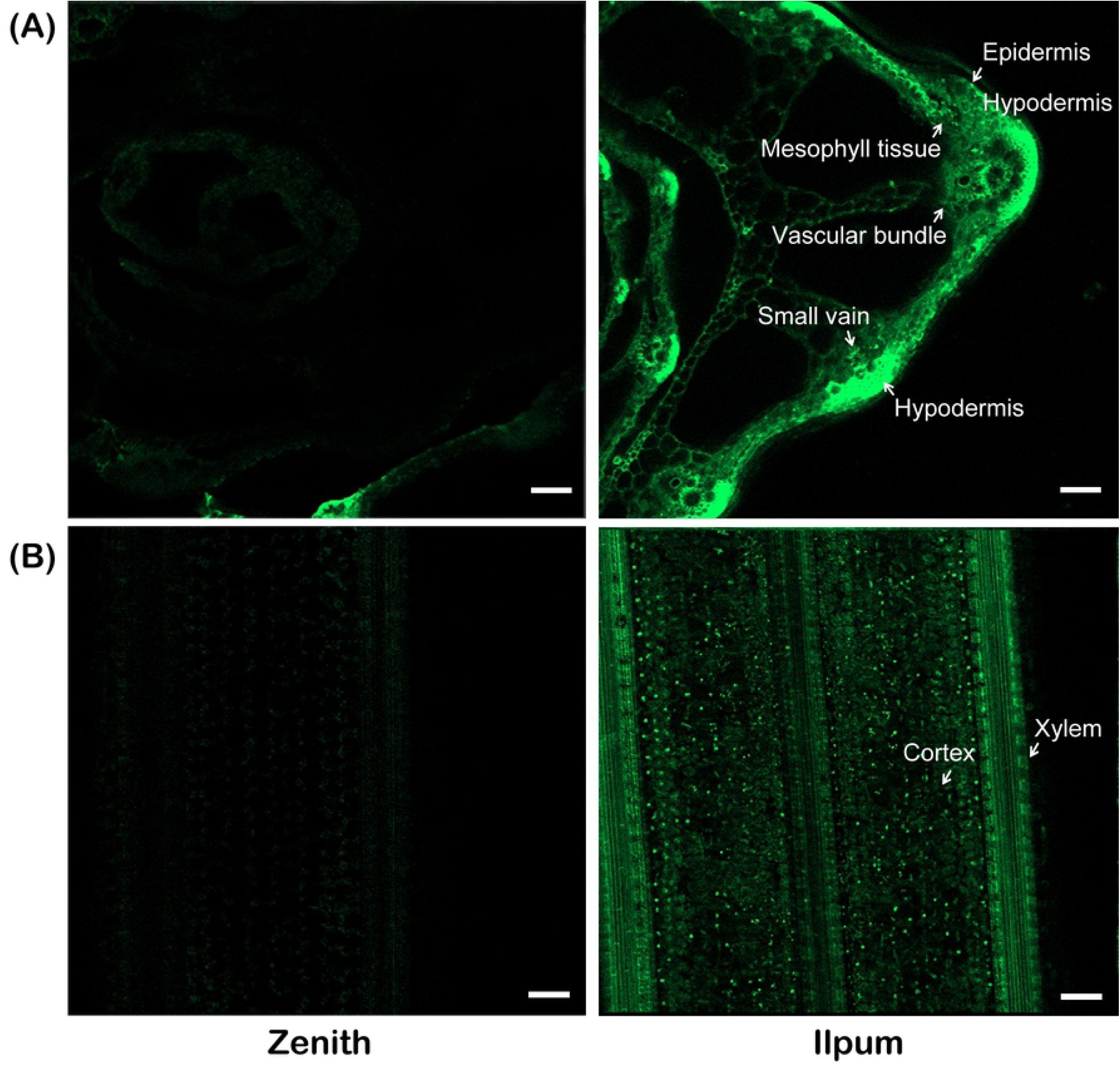
Confocal imaging of shoot base of 10 days-old-seedling derived from Zenith and Ilpum rice plants infected with *F. fujikuroi* (CF283 isolates) containing GFP expression cassette. (A) Radial and longitudinal sections of the basal stem (Scale bar = 50 μm).

### QTL analysis and mapping of *qBK1*^*z*^ using 180 F_2:9_ RILs

Based on the bakanae disease bioassay (proportion of healthy plants), the 180 F_2:9_ RIL population exhibited continuous distribution (0.0–98.0%; Fig. 3), which quantitatively confirmed the inheritance of bakanae disease resistance. The average proportion of healthy Ilpum plants was 14.3% (3.3–37.0%; *n* = 10) and that of Zenith was 63.2% (42.7–79.3%; *n* = 10).

**Fig 3.**
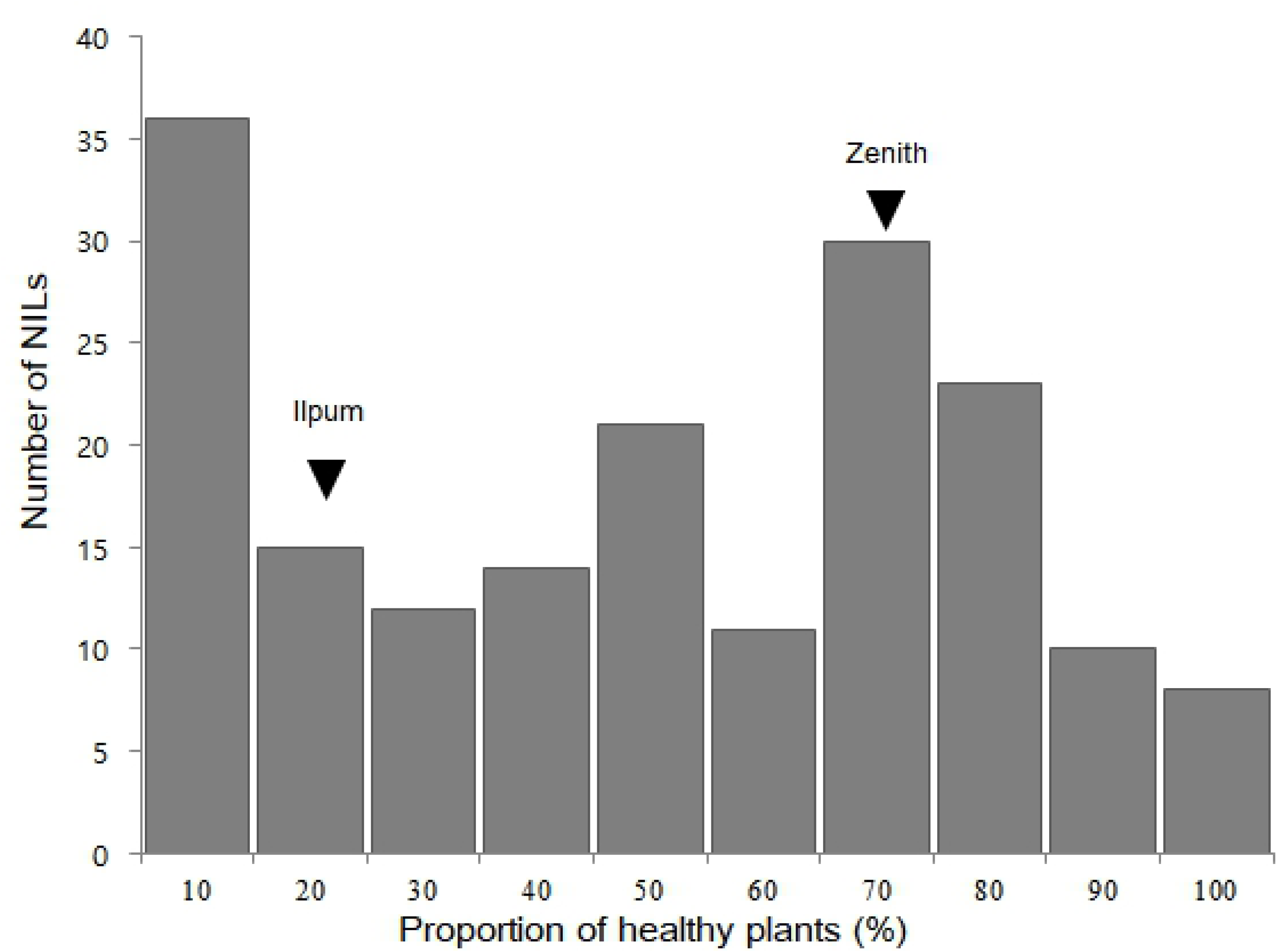
Frequency distribution of the proportion of healthy F_2:9_ NILs derived from a cross between Ilpum and Zenith after bakanae disease inoculation. The mean proportion of healthy plants of Ilpum and Zenith is indicated by black arrow heads.

We selected 164 SSR markers among 1,150 SSR markers from the polymorphism survey between Ilpum and Zenith from the Gramene database (http://www.gramene.org) covering the whole rice chromosome (S1 Fig). The genetic linkage map of Ilpum and Zenith for primary mapping was constructed with 164 polymorphic markers covering a total length of 3,140 cM with average interval of 19.14 cM as described by Lee et al. [30]. Primary QTL mapping using the 180 F_2:9_ populations showed that a significant QTL associated with bakanae disease resistance at the seedling stage was located between the SSR markers, RM1331 and RM3530 on chromosome 1, and it was designated *qBK1*^*z*^. The LOD score of *qBK1*^*z*^ was 13.43, which accounted for 30.9% of the total phenotypic variation (Table 1).

**Table 1.**
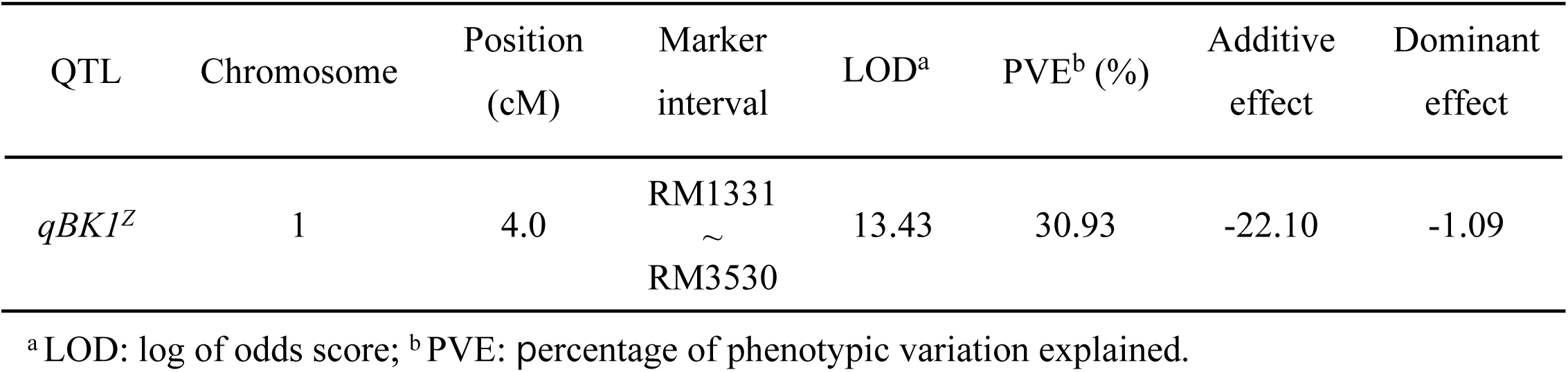
Putative quantitative trait locus (QTL) associated with bakanae disease resistance detected at the seedling stage by composite interval mapping of the 180 F_2:9_ populations derived from a cross between Ilpum and Zenith

A finer localization of *qBK1*^*z*^ was determined by analyzing the chromosome segment introgression lines in the region detected from primary mapping. The *qBK1*^*z*^ region between RM1331 and RM3530 from primary mapping was narrow downed with additional 55 SSR markers and designed 12 InDel markers for the insertion/deletion sites based on the differences between the *japonica* (http://www.gramene.org) and *indica* (http://rice.genomics.org.cn) sequences. Four SSR markers and six InDel markers were selected as polymorphic markers between the parents to narrow down the position of the *qBK1*^*z*^ region (S2 Table). Finally, seven homozygous recombinants were selected from the F_2:9_ lines using 14 markers in the 2.8 Mb region around the SSR markers RM1331 and RM3530 (Fig. 4 and Fig. 5).

**Fig 4.**
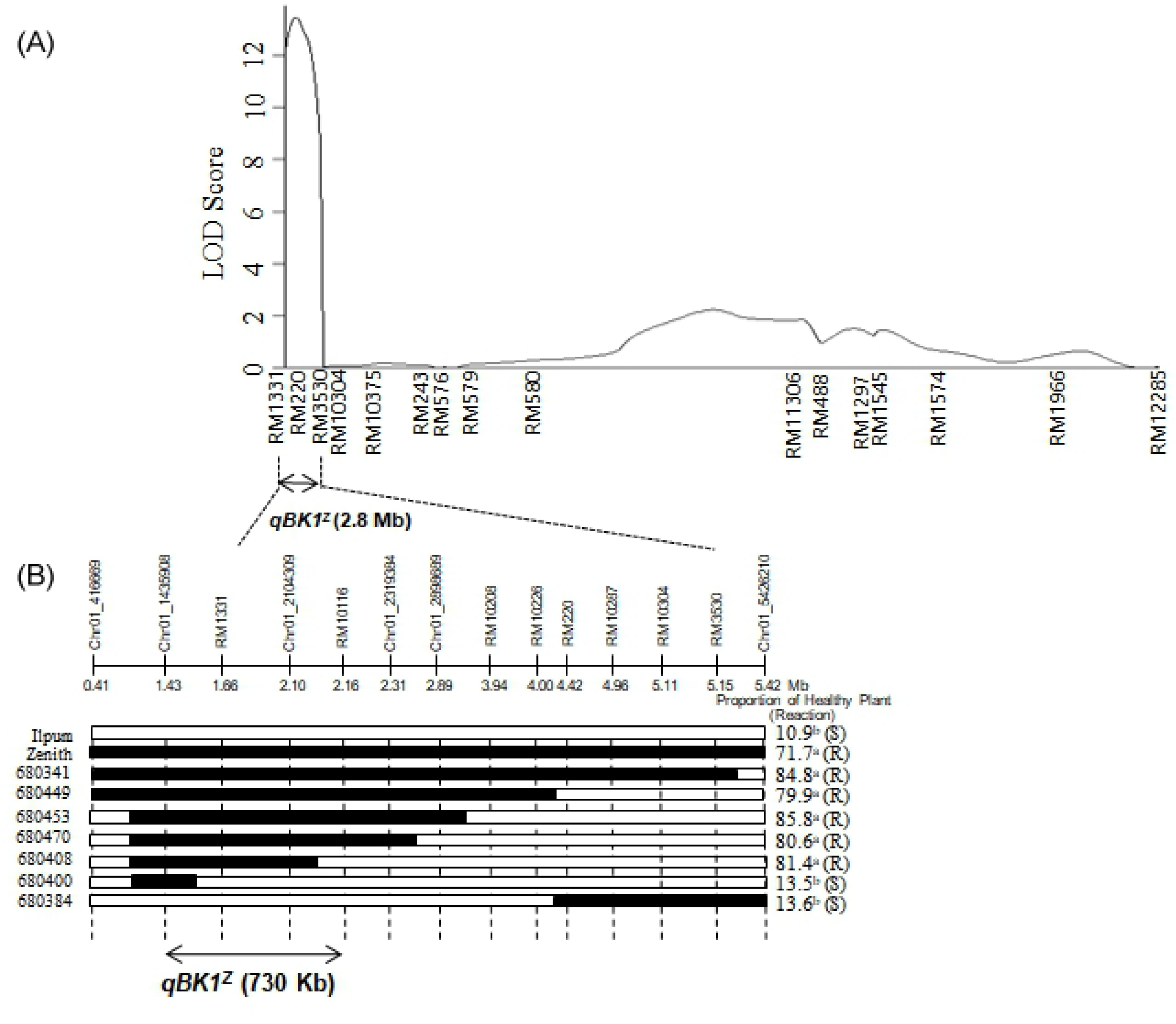
QTL analysis of *qBK1*^*Z*^ on chromosome 1 using 180 RILs derived from a cross between Ilpum and Zenith. (a) In primary mapping, *qBK1*^*Z*^ was located in 2.8 Mb regions between the RM1331 and RM3530 markers on chromosome 1. (b) Location of *qBK1*^*Z*^ was narrowed down to 730 kb region between the markers Chr01_1435908 and RM10116 in secondary mapping using selected seven homozygous recombinants. Black bars show the homozygous regions for zenith alleles; white bars indicate the homozygous regions for Ilpum alleles. R; resistant to bakanae disease, S; susceptible to bakanae disease. The proportion of healthy plant was calculated from three biological replications. Values (%) of the on proportion of healthy plant with different letters are significantly different by Duncan’s multiple range test at P = 0.05.

**Fig 5.**
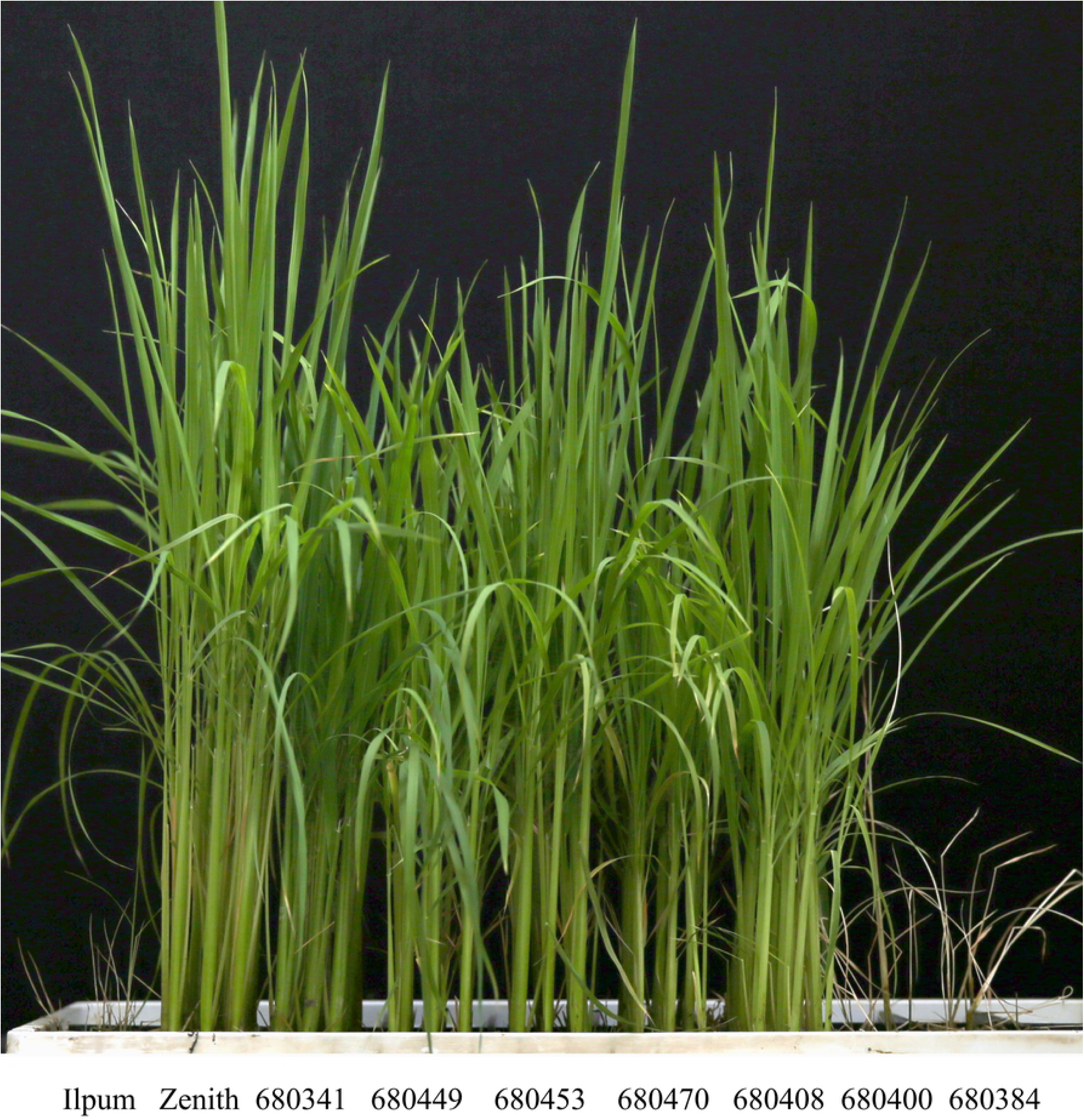
Phenotypic responses to bakanae disease infection of seven homozygous recombinants used in secondary mapping.

The proportion of healthy plants of the seven homozygous recombinants was evaluated with three biological replicates according to Duncan’s new multiple range test. Based on this bioassay, lines classified to Group a were regarded as resistant, and Group b as susceptible (Fig. 4). Considering the genotype and the phenotype of the recombinants, it is clear that *qSTV11*^*Z*^ conferring resistance to bakanae was an approximate 730 kb interval delimited by the physical map between Chr01_1435908 (1.43 Mbp) and RM10116 (2.16 Mbp).

## Discussion

Rice varieties with a single resistance gene are at an increased risk of being overcome by new pathological races [15, 30]. The development of a rice variety with a higher level of resistance against bakanae disease is a major challenge in many countries [9, 20, 31-33]. In this study, we identified *qBK1*^*z*^ locus related to bakanae disease resistance based on genotype and phenotype analyses of homozygous recombinants on the recombinant progeny of Ilpum and Zenith, using SSR and newly developed InDel markers.

Localization of *F. fujikuroi* isolate CF283 in the susceptible Ilpum and the resistant Zenith was also examined. It was reported that successful infection of *Fusarium* species is a complex process that includes adhesion, penetration (through wounds, seeds, stomatal pores) and subsequent colonization within and between cells [34, 35]. Consistent with previous reports [36, 37], *F. fujikuroi* isolate CF283 was extensively observed on the aerenchyma, pith, cortex and vascular bundle of both leaf and stem in susceptible Ilpum, whereas this was rarely observed in resistant Zenith.

Many QTLs on bakanae disease resistance have been identified on chromosome 1. Three QTLs, *qBK1*^*z*^, *qBK1*.*2 and qBK1*.*3*, were found in a similar region in spite of the different source of resistant varieties (Fig 6).

**Fig 6.**
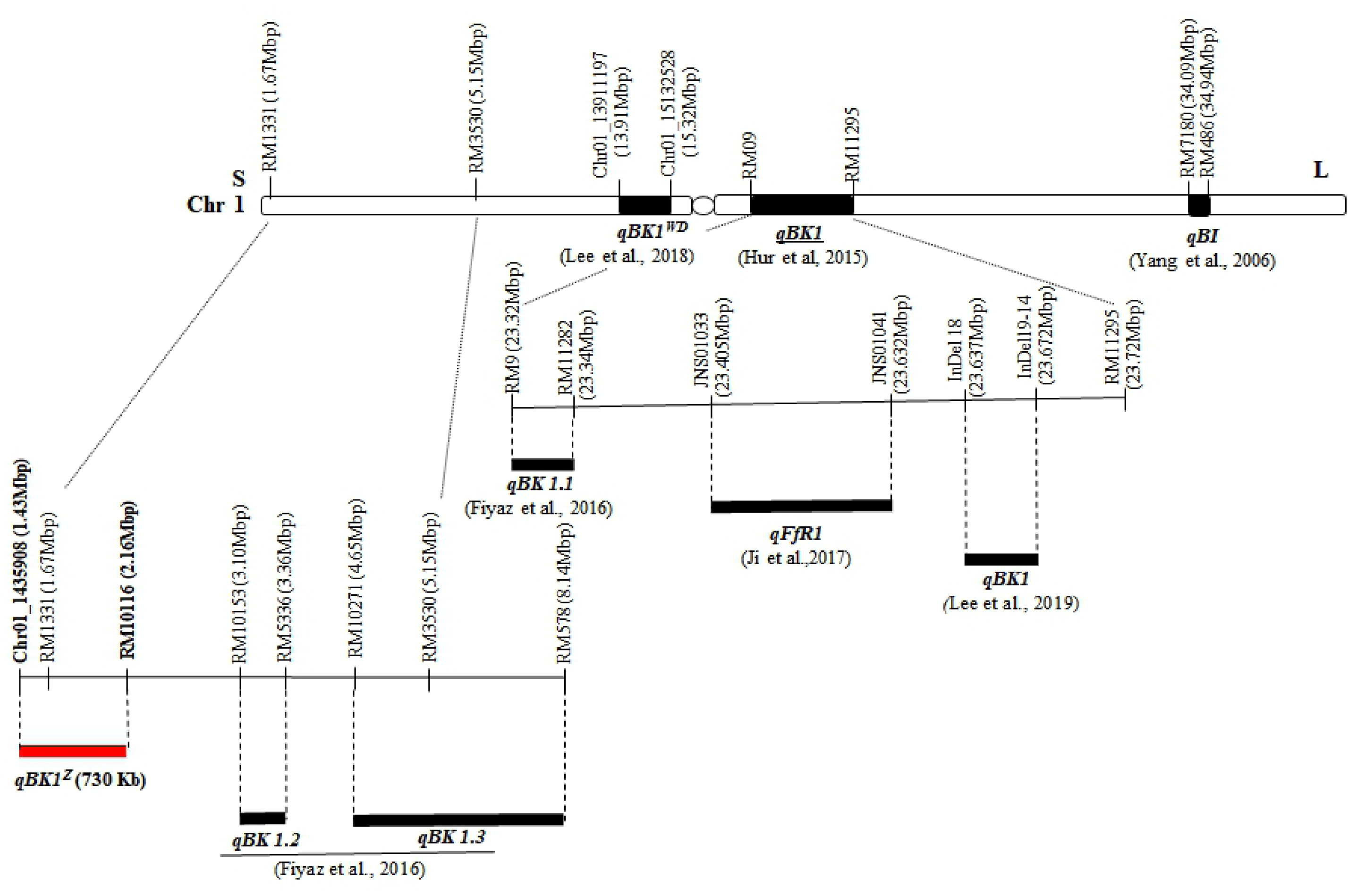
Physical locations of bakanae disease resistance QTLs identified on chromosome 1.

Fiyaz et al. [9] mapped *qBK1*.*2* to a 260 kb region between markers RM10153 and RM5336, and *qBK1*.*3* to a 3.49 Mb region between markers RM10271 and RM578 from the Pusa 1121/Pusa1342 cross. Location of *qBK1*^*z*^ was found upstream to *qBK1*.*2*. Fiyaz et al. (2016) mapped *qBK1*.*1* to a 20 kb region between markers RM9 and RM11232 from the Pusa 1121/Pusa1342 cross. These authors hypothesized that *qBK1*.*1* and *qBK1* [20] might be the same QTL as they had overlapping positions. Ji et al. [22] found that QTL *qFfR1* was located in a 230 kb region of rice chromosome 1 in Korean *japonica* variety Nampyeong, and suggested that the three QTLs *qBK1, qBK1*.*1*, and *qFfR1* might indicate the same gene. Lee et al. [21] narrowed down the position of the *qBK1* locus to a 35 kb region between InDel 18 and InDel 19-14, and revealed that location of *qBK1* is close to those of *qBK1*.*1*, and *qFfR1*, does not overlapping each other. Two additional QTLs including *qB1* from Chunjiang 06 [19] and *qBK1*^*WD*^ from Wonseadaesoo [15] also were also found on chromosome 1. Gene pyramiding via phenotypic screening assays for crop breeding is considered to be difficult and often impossible due to dominance and epistatic effects of genes governing disease resistance, and the limitation of screenings being all year-round [38]. Pyramiding of multiple resistant QTLs/genes by using marker-assisted breeding (MAB) in a single plant might confer higher and/or durable resistance against bakanae disease. The effects of pyramiding resistance genes have been observed for several plant-microbe interactions. Pyramiding three bacterial blight resistance genes resulted in a high level of resistance and were expected to provide a durable pathogen resistance [39, 40]. On the other hand, pyramiding of resistant genes resulted in a level of resistance that was comparable to or even lower to than that of the line with a single gene. For example, Yasuda et al. [41] reported rice lines with pairs of blast resistance genes to be only comparable to lines with a single gene which may have a stronger suppressive effect. Similarly, pyramiding of borwn planthopper (BPH) resistance genes in a susceptible variety IR24 markedly improved BPH resistance against four BPH populations in the Philippines than the single BPH introgressed lines [42].

In our previous study of bakanae disease resistance [15] revealed that the lines harboring *qBK1*^*WD*^ showed a higher resistance than those with *qBK1*. Furthermore, the pyramided lines harboring *qBK1*^*WD*^ + *qBK1* had a much higher level of resistance than those possessing either *qBK1*^*WD*^ or *qBK1*. These results showed the utility of MAB in gene pyramiding can achieve higher resistance in many bakanae disease prone rice growing areas.

## Conclusions

In this study, we identified a new major QTL *qBK1*^*z*^ conferring bakanae disease resistance from a new genetic source of *indica* variety, Zenith. Through QTL analysis and fine mapping, we narrowed down the *qBK1*^*z*^ locus into 730 kb on the short arm of chromosome 1 where is a novel locus compared with all the previously identified bakane disease resistance QTLs. Together with the previously identified QTLs of the bakane disease resistance, the new *qBK1*^*z*^ can be introduce to the elite favorable background varieties by aa marker-assisted backcrossed breeding. Furthermore, the new QTL *qBK1*^*z*^ will be a useful material for studying an interaction between the pathogen (*F. fujikuroi*) and rice host plants.

## Supporting information

S1 Fig. Linkage map constructed with 180 F_2:9_ recombinant inbred lines (RILs) derived from a cross between Zenith and Ilpum. (PPT)

S2 Table. InDel and SSR markers used for the fine mapping of *qBK*^*z*^. (DOCX)

## Author contributions

Conceptualization: SB Lee, YJ Hur, DS Park

Data curation: SB Lee, N Kim, YS Seo, J Lee, DS Park

Formal analysis: SB Lee, DS Park

Funding acquisition: DS Park

Investigation: SB Lee, N Kim

Methodology: SB Lee, DS Park

Project administration: JM Ko, DS Park

Software: SM Jo, JH Cho

Supervision: DS Park

Validation: YS Seo, J Lee, DS Park

Visualization: JHee Lee, J Lee, JW Kang, YC Song.

Writing – original draft: SB Lee, DS Park

Writing – review & editing: YS Seo, J Lee, M Bombay, SR Kim, DS Park

